# MOSAIC: A Pipeline for MicrobiOme Studies Analytical Integration and Correction

**DOI:** 10.1101/2024.11.07.622561

**Authors:** Chenlian Fu, Jiuyao Lu, Ni Zhao, Wodan Ling

## Abstract

Large-scale and consortium microbiome studies have enabled identification of reliable population-level biomedical signals, wherein integration is essential to eliminate unwanted variations between batches or studies and retain biological signals. Many strategies, each with distinct advantages and limitations, have been adapted or developed for microbiome data. The optimal strategy for a given study needs to be determined on a data-specific, case-by-case basis. Here, we develop the first-of-its-kind MicrobiOme Studies Analytical Integration and Correction (MOSAIC) pipeline to enable a convenient, fair, and comprehensive comparison of integration strategies. It includes modules for pre-processing, integration, and evaluation of artifact removal and signal preservation, using metrics relevant to common microbiome analyses, including alpha and beta diversities, disease prediction, and differential abundance analysis. We applied MOSAIC to extensive real-world and simulated data and found that though no single strategy excels in all aspects, yet certain strategies, the ComBat and ConQuR families, perform better overall.

## Introduction

The microbiome has been shown to play critical roles in human health and disease, such as type II diabetes (T2D) ^1–3^, inflammatory bowel disease (IBD) ^4–7^, colorectal cancer (CRC) ^8–11^, among others. Given the importance of microbiome and the reduced cost of profiling technologies, large-scale and consortium microbiome studies, which involve multiple batches or studies on the same disease, have emerged and become the trend ^5,12,13^. Analyzing the corresponding microbiome data, which is combined from different sources, is termed horizontal data integration ^14^. The horizontal integration enables investigators to achieve population-level discoveries, helping uncover signals that are unattainable with the sample size of a single batch or study (improving statistical power) and avoiding spurious findings caused by biases in individual batches or studies (enhancing statistical robustness).

Despite these advantages, batch effect or between-study heterogeneity pose a critical challenge in the integrative analyses. In large-scale studies, samples have to be collected and processed in separate runs, introducing variations from differential handling and processing of specimens^15^. Consortium studies add even more heterogeneity due to varying study designs, experimental protocols, and profiling technologies ^16^. In this paper, we refer to these unwanted variations, whether due to batches or studies, as batch effects. Failing to mitigate batch effects can obscure true biological signals and lead to spurious findings. Thereafter, many integration strategies have been adapted or developed for microbiome data, aiming to remove batch effects while preserving the biological signals of interest.

Commonly used strategies that are adapted from other omics field include the ComBat ^17^ family (ComBat and ComBat-seq ^18^), limma ^19^ (the removeBatchEffect module), and Harmony ^4^. ComBat ^17^ and limma ^19^ are popular approaches designed for gene expression data, with the former primarily designed for microarray data and latter for both microarray and bulk RNA-seq expression data. ComBat-seq ^18^ is an extension of ComBat ^2^ to bulk RNA-seq count data. Harmony was developed for scRNA-seq data. It identifies groups of cells with similar expression patterns across batches (called anchors), and then uses similarity-based methods to align these similar cells to a common distribution in a reduced latent space, regardless of batch. Tailored strategies for microbiome data have been developed in recent years, such as the ConQuR family ^20^, MMUPHin ^21^, and Percentile Normalization ^22^. Each strategy makes its own assumptions (or assumption-free) of microbiome data and models it accordingly to remove batch effects. Though these strategies have demonstrated success in mitigating batch effects while improving biomarker discovery and phenotype prediction in their own publications ^21,23,24^, their performance may not be universal across different data sets. The optimal integration strategy depends on the specific characteristics of a given data, such as the extent of sparsity and over-dispersion, which are influenced by the body site, study design, profiling technologies, etc. Consequently, selecting the best integration strategy should be done on a case-by-case basis. However, this selection process is challenging. Implementing all strategies while considering their unique requirements and configurations (Supplemental Table 1) is tedious, and there is no systematic framework to evaluate the effects of various strategies. These gaps necessitate a computational pipeline that allows researchers to easily implement and choose the optimal integration strategy for their data and analytical goals through a fair and comprehensive comparison.

Here, we present a first-of-its-kind analytical pipeline, MOSAIC (MicrobiOme Studies Analytical Integration and Correction), which streamlines the implementation and evaluation of various integration strategies for microbiome data. MOSAIC manages the implementation details of each strategy, ensuring optimal use and fair comparison. After applying these integration strategies, MOSAIC provides a comprehensive assessment of the integrated data, focusing on their effectiveness in removing batch effects and preserving key variable effects. This quantitative evaluation caters to a wide range of microbiome analyses, including alpha and beta diversities, microbiome-based predictions for the key variable, and differential abundance analysis. Facilitated by MOSAIC, we compare the integration strategies across extensive real-world and simulated microbiome studies, encompassing multiple diseases, varied degrees of batch and key variable effects, different types of the original microbiome data (taxonomic count or relative abundance), etc. The final reports from MOSAIC reveal that no method can dominate at all evaluation criteria, while some strategies can perform better overall, such as the ComBat and ConQuR families.

## Results

### MOSAIC: a pipeline for systematic implementation and comparison of microbiome integration strategies

The goal of MOSAIC is to enable a convenient, fair, and comprehensive comparison of various integration strategies for microbiome studies. To achieve this, MOSAIC is divided into three modules: pre-processing, integration, and evaluation. Initially, MOSAIC pre-processes the microbiome and metadata, conducting taxonomic aggregation, filtering of rare taxa, and removal of low-quality sample, as well as fulfilling specific requirements for each integration strategy (Supplemental Table 1). The processed data are fed into the corresponding integration strategies to generate integrated and batch-corrected data. Finally, the pipeline evaluates the effectiveness of each strategy in removing batch effects and preserving key variable effects.

A wide range of metrics, inspired by four common microbiome analytical objectives, are used in the evaluation. The first two metrics are based on alpha and beta diversities. Alpha diversity (e.g. Shannon index) reflects richness and/or evenness of each sample’s microbial profile. An effective integration strategy reduces the association between the batch and alpha diversity while strengthening the association between the condition and alpha diversity. We therefore assess this association by statistical significance. Beta diversity, encoded by ecologically informative distance metrics such as Bray-Curtis and Aitchison dissimilarities, allows us to evaluate the proportion of variability in microbiome composition explained by the batch or condition, quantified by PERMANOVA ^25^ R^2^. A successful integration is shown by a reduced batch R^2^ with a maintained or increased condition R^2^. The third metric evaluates the explanatory capability of microbial profiles for the key variable of interest. The fourth metric relates to differential abundance analysis, identifying microbial taxa associated with the key condition. False discovery rate (FDR) and sensitivity are used to show how integration improves the reliability of biomarker discovery. Details of the metrics can be found in the Methods section.

In summary, MOSAIC facilitates fair comparisons among them through a comprehensive suite of evaluation metrics (Figure 1) to help researchers select strategies best fitting their analytical needs. MOSAIC is packaged into a Snakemake ^26^ pipeline, allowing easy instruction customization and execution via a simple command line.

**Figure 1.**
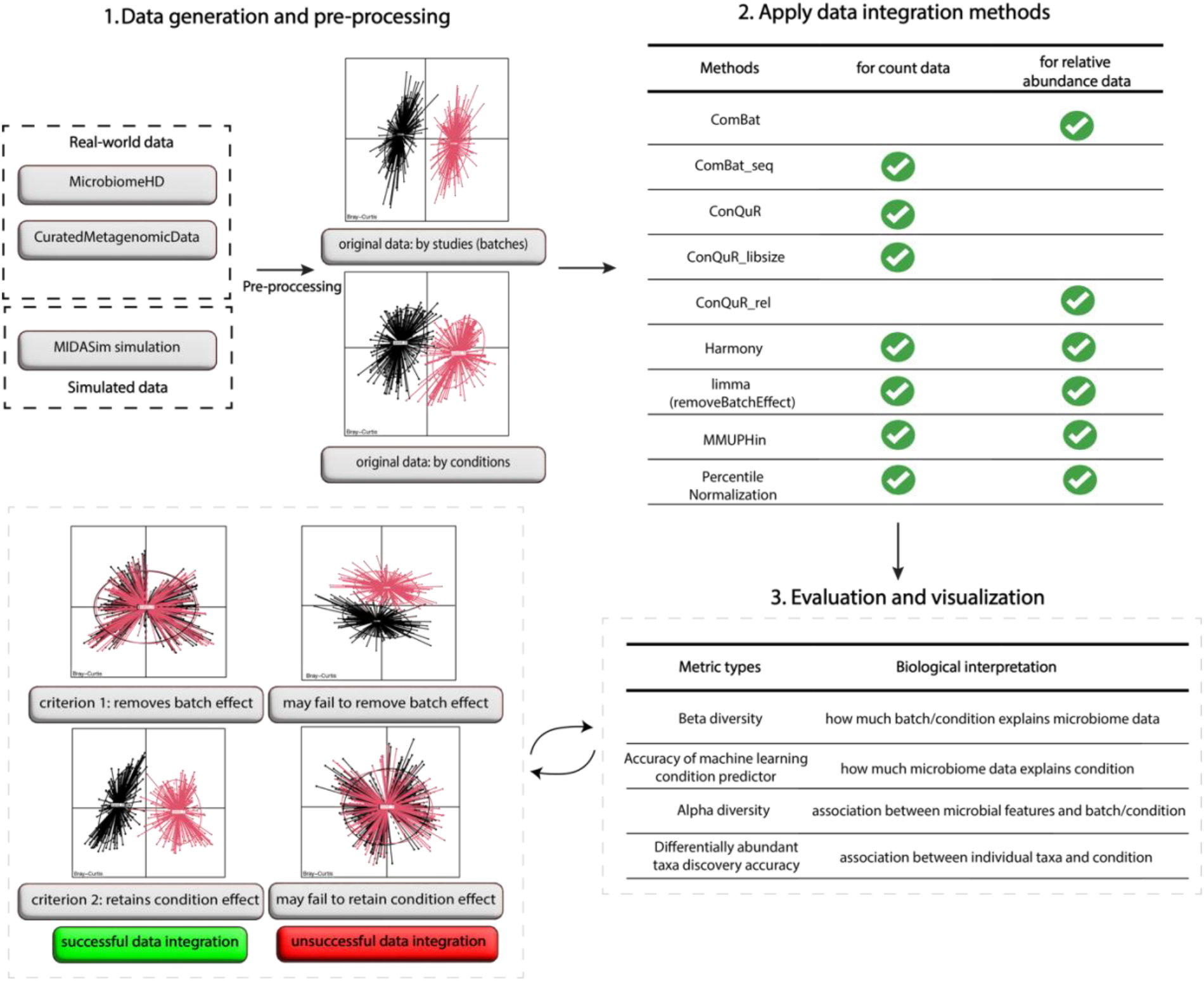
Graphic summary of MOSAIC. First, real-world data or simulated data go through the respective pre-processing steps. Then, the processed data undergo the respective data integration strategies according to the nature of the processed data (taxonomic count or relative abundance). The pipeline then provides a comprehensive evaluation and visualization to compare different integration strategies’ effectiveness according to two criteria – 1) remove batch effect and 2) retain the condition effect – using multiple metrics inspired by common microbiome analyses.

### Evaluating integration strategies on simulated microbiome studies via MOSAIC

We simulated data using MIDASim ^27^ to resemble a cleaned, real gut microbiome IBD data set from HMP2Data ^28^ (details in Methods). Specifically, we simulated two conditions and two batches with varying levels of confounding effects between them. A wide spectrum of batch and condition effects were considered, including null case (batch effect=0), and when batch effect is bigger than, smaller than, or equal to the condition effect. Additionally, we varied whether library size difference is part of the batch effect. Both the simulated taxonomic count data and the corresponding relative abundance data are considered in evaluating integration strategies.

In simulated taxonomic count data, ComBat-seq and ConQuR demonstrated overall satisfactory performance in both batch effect removal and condition effect preservation.

Regarding the association between microbial richness and evenness and batch/condition (Figure 2b, left/right), at a significance level of = 0.05, Percentile Normalization thoroughly removed batch effects (controlled type I error for batch around 0.05) but also eliminated condition effects (reduced power for condition to 0.05). ConQuR was the second most effective in removing batch effects (controlled type I error for batch below 0.05, except in one scenario with the largest batch effect). Simultaneously, it maintained the condition effect, achieving the highest power for condition among all strategies. ComBat-seq was generally effective, but its performance in both batch effect removal and condition effect maintenance diminished when the condition effect was large. The remaining strategies failed to remove batch effects (kept type I error for batch above 0.05). Among them, MMUPHin retained the condition effect, ConQuR-libsize and Harmony preserved partial condition effect, while limma almost eliminated the condition effect.

**Figure 2.**
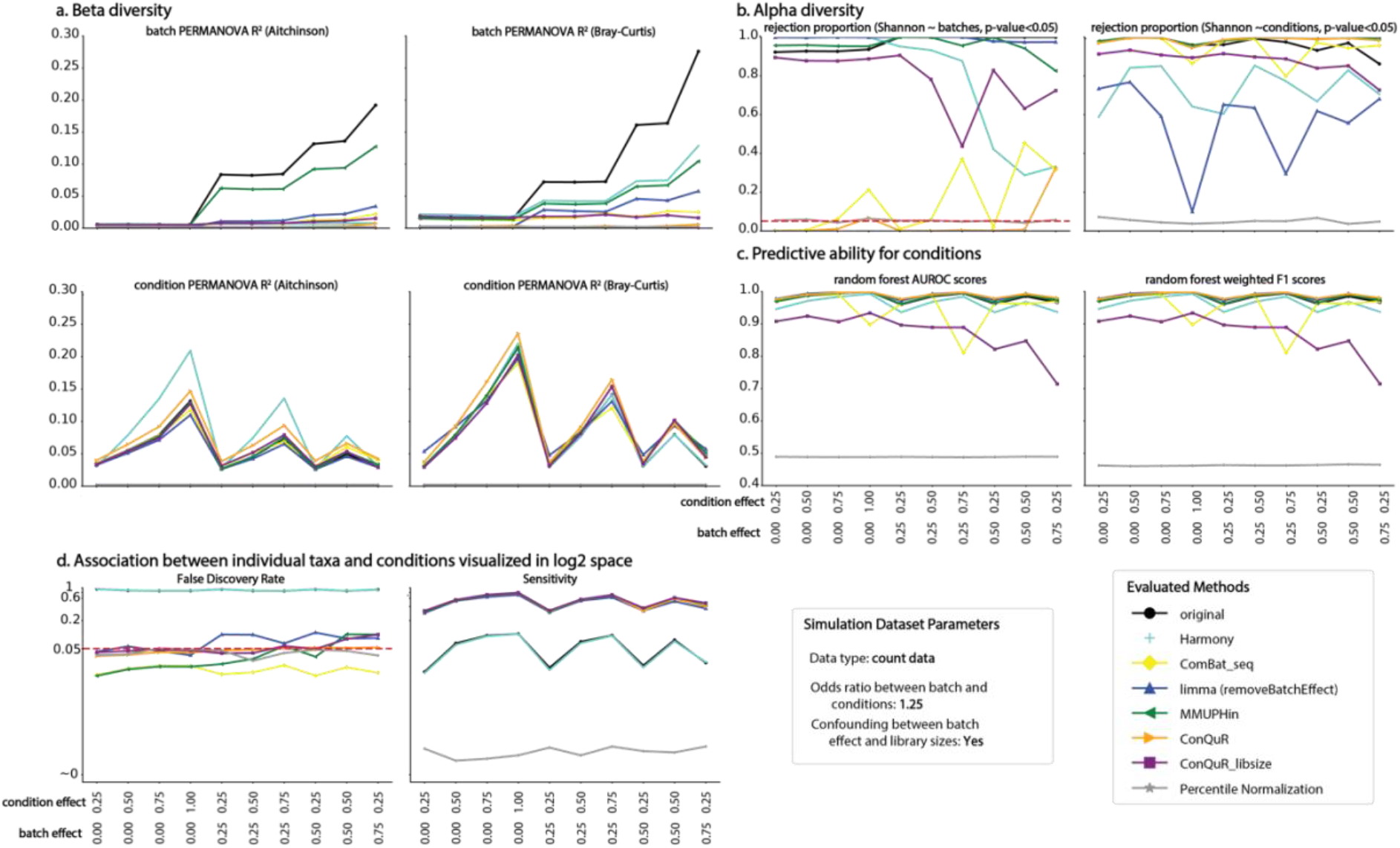
Evaluation of data integration strategies on simulated microbiome data. Taxonomic count data were generated by MIDASim to mimic the HMP-IBD data with 2 conditions and 2 batches. Condition and Batch are simulated from a joint Bernoulli *distribution with p*_*Condition*_=0.5, *p*_*batch*_=0.5, and OR=1.25. A wide range of condition and batch effects were examined, covering batch effect < condition effect, batch effect= condition effect, and batch effect > condition effect. For each panel, the scenarios are arranged by the increasing batch effect and the increasing condition effect within each value of batch effect. a. Proportion of microbiome variability explained by batch and condition, as measured by the average PERMANOVA R^2^ in Aitchison and Bray-Curtis dissimilarities. b. Rejection proportion of association significance between alpha diversity (Shannon index) and batch/condition (i.e., type I error/power) using Wilcoxon rank-sum test at α= 0.05. c. Average cross-validated prediction accuracy (AUROC and F1) of the integrated microbiome data for the condition using random forest. d. Average FDR and sensitivity of individual taxa’s association with the condition (visualized in log2 space).

Regarding the variability in microbiome composition explained by batch/condition (Figure 2a, top/bottom), Percentile Normalization removed the most batch effects but also erased condition signals altogether. ConQuR performed equivalently to Percentile Normalization in removing batch effects but retained or even amplified the condition effect. ConQuR’s efficacy was followed by ComBat-seq and ConQuR-libsize, although these methods could not thoroughly remove batch effects when library size differences contributed to batch heterogeneity (Figure 2a, batch PERMANOVA R2 in Bray-Curtis). Harmony, limma, and MMUPHin, exhibited comparable effectiveness in partially reducing batch effects. However, MMUPHin and Harmony were among the top tier in retaining condition signals.

The predictive metric allowed us to further assess how effectively the microbiome could explain the condition effect. Figure 2c shows that the Percentile Normalization-integrated microbial profile no longer possessed condition signals (AUROC around 0.5, equivalent to random guessing). Most strategies, including ConQuR, MMUPHin, Harmony, and limma, performed comparably well in retaining the condition effect. However, ComBat-seq diluted the condition effect when the simulated condition signals were strong, and ConQuR-libsize was generally worse-performing, especially when batch effect was large.

Lastly, in differential abundance analysis, almost all strategies, except for Harmony, maintained FDR around 0.05. However, MMUPHin and ConQuR-libsize showed noticeably inflated FDR under large batch effect. Among the strategies that generally controlled FDR, MMUPHin, ConQuR, ConQuR-libsize, and ComBat-seq improved the power of differential taxa identification, enabling reliable and informative biomarker discovery.

Under settings with varied confounding effects between batch and condition and varied confounding statuses between batch and library size, as well as in relative abundance data (Supplemental Figures 1-4), the comparison results among the investigated strategies, in general, remain consistent.

In summary, different integration methods excel at different aspects of microbiome data analysis. Percentile Normalization excels in batch removal across the board but maintains minimal biological signals for downstream analyses. Most other methods excel at microbiome-based phenotype prediction and individual taxa identification, with ComBat-seq, ConQuR family (for respective data types), MMUPHin particularly outstanding. Alpha and beta diversity metrics show nuanced batch/condition effect trade-offs in different strategies and highlight the robustness of ConQuR and ComBat families. It is also worth noting that computational time is critical, especially when resources are limited and integration is just one of the upstream analysis steps. The average runtime across 1,000 simulation runs shows that the ConQuR family is consistently slower than their counterparts (Supplemental Figure 5a-c).

### Evaluating integration strategies on real-world microbiome studies via MOSAIC

Facilitated by MOSAIC, we benchmarked integration strategies for four real-world microbiome studies (Supplemental Table 1, 2). Two of them are genus-level count datasets generated using used 16S rRNA sequencing from the MicrobiomeHD database ^24^, focusing on autism and Clostridioides difficile infection (CDI) patients and their controls, and include two and three sub-studies, respectively. For the other two studies, we prepared relative abundance data at the species level generated from shotgun metagenomic sequencing from the CuratedMetagenomicData database ^29^, pertaining to IBD and CRC patients and their controls, and include three and eight sub-studies. Disease status such as autism vs. control is the key condition of interest.

For the autism and CDI studies with count data, all strategies reduced batch effects compared to the data before integration, with Percentile Normalization, ConQuR, and limma being the most outstanding (Figure 3a-b). Regarding the condition effect (Figure 3b), the autism data represents a real-world scenario where condition effect is minimal prior to integration, leaving a limited room for signal retainment; for the CDI data, ComBat-seq and MMUPHin, followed by limma, ConQuR, and ConQuR-libsize, demonstrated relative effectiveness in retaining condition effects. Considering the richness/evenness of the microbial profiles, Percentile Normalization and ConQuR were the only strategies capable of making batch not associated with the characteristics (Figure 3c, p-value > 0.05) in both the autism and CDI examples. However, no methods can improve the association between the richness/evenness and autism status, while only Percentile Normalization failed to maintain the association between the richness/evenness and CDI status. Regarding predictive abilities for disease statuses, ConQuR and ConQuR-libsize improved condition signals, while the others maintained or even lost the key signals.

**Figure 3.**
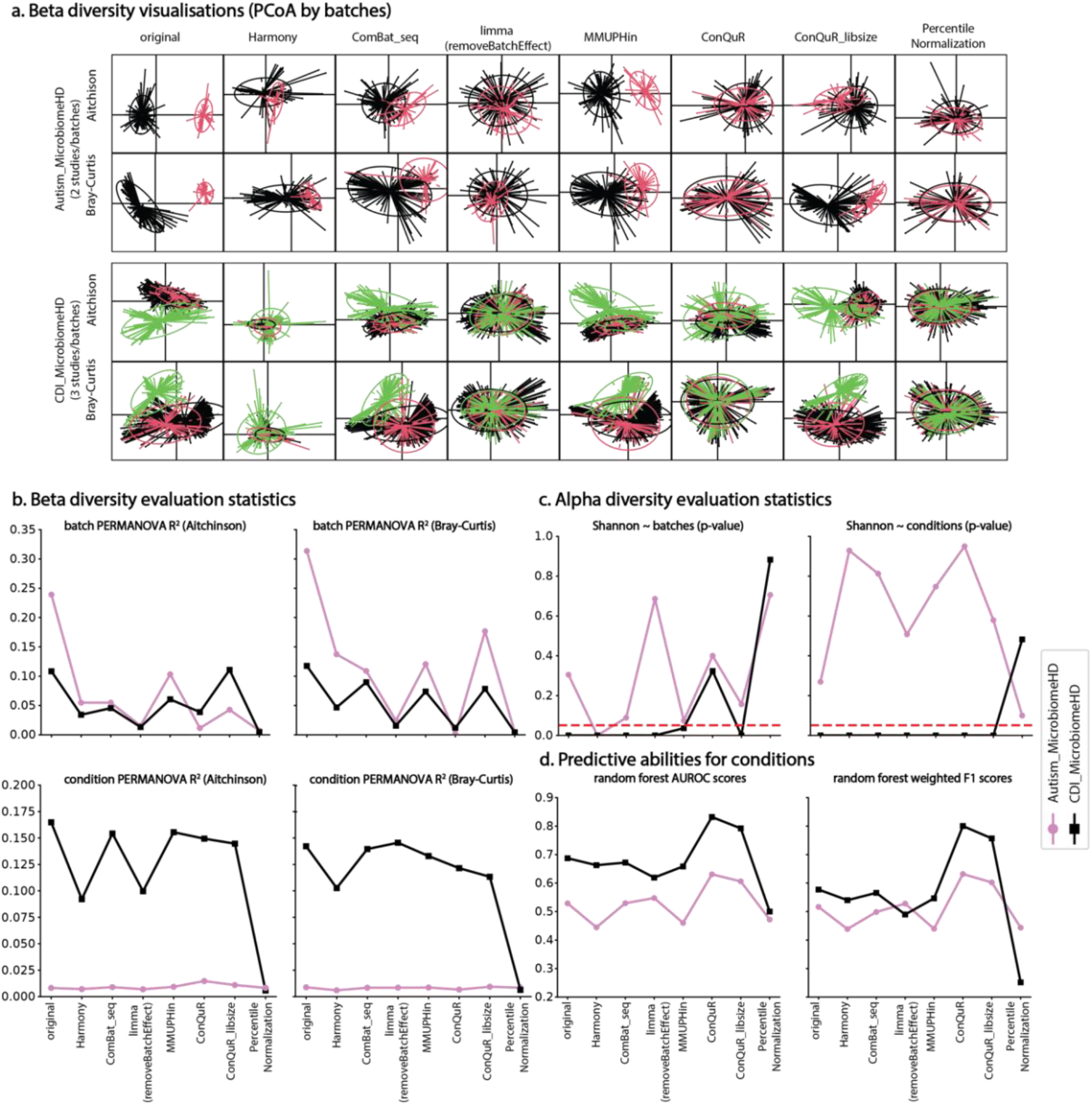
Evaluation of data integration strategies on real-world microbiome data of taxonomic count. Showed with autism data with 2 studies and CDI data with 3 studies. a. PCoA plots in Aitchison and Bray-Curtis dissimilarities stratified by studies. b. Microbiome variability explained by study and disease condition, as measured by the PERMANOVA R^2^ in Aitchison and Bray-Curtis dissimilarities. A smaller study R^2^ and a larger condition R^2^ are preferred. c. Association significance (p-value) between alpha diversity (Shannon index) and batch/condition using Wilcoxon rank-sum test at α= 0.05. A non-significant association between alpha diversity and batch and a significant association between alpha diversity and condition are preferred. d. Cross-validated prediction accuracy (AUROC and F1) of the integrated microbiome data for the disease condition using random forest. Higher AUROC and F1 indicate a better prediction combining sensitivity and specificity.

**Figure 4.**
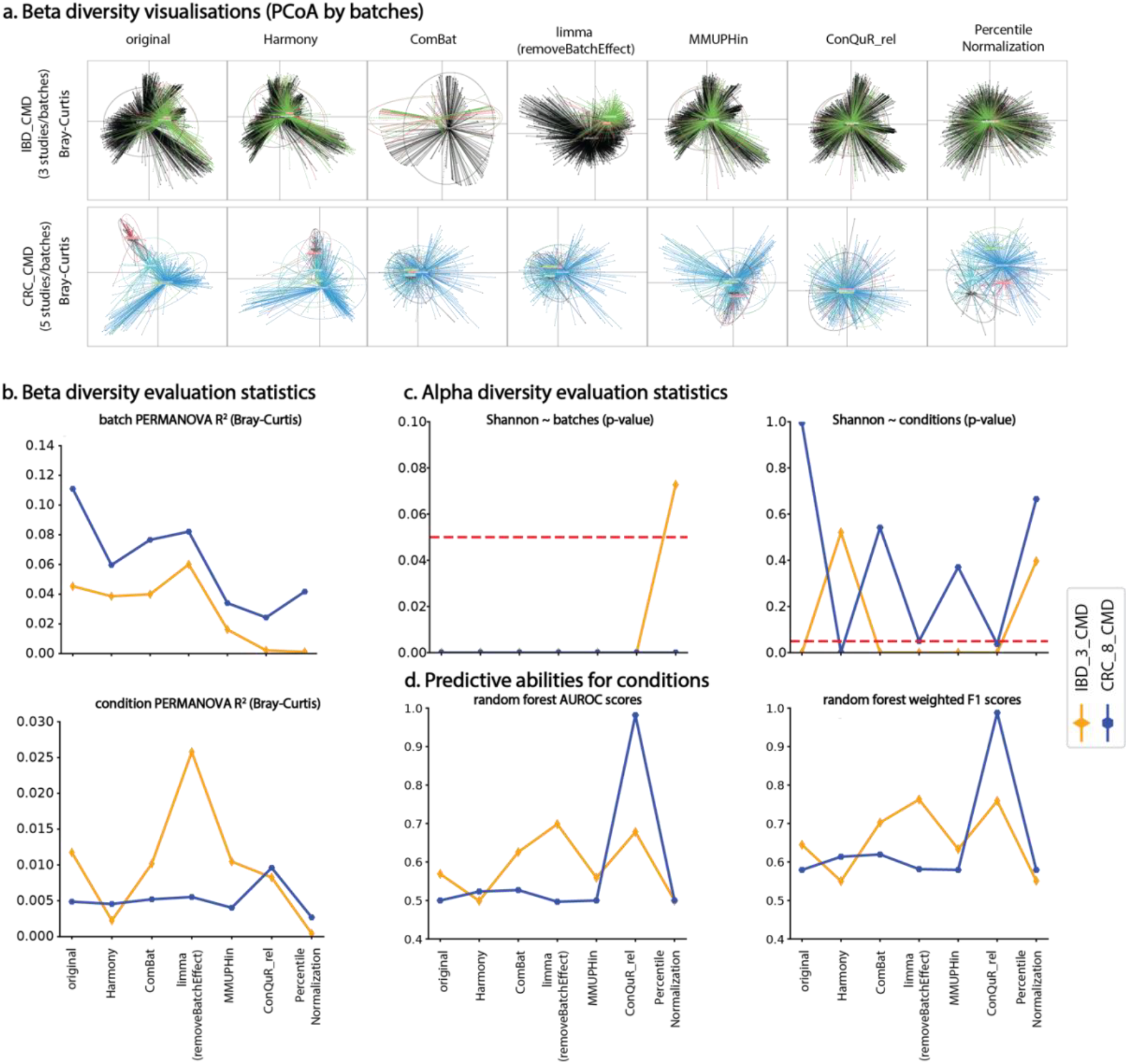
Evaluation of data integration strategies on real-world microbiome data of relative abundance. Showed with IBD data with 3 studies and CRC data with 5 studies. a. PCoA plots in Bray-Curtis dissimilarity stratified by the studies. b. Microbiome variability explained by study and disease condition, as measured by the PERMANOVA R^2^ in Bray-Curtis dissimilarity. A smaller study R^2^ and a larger condition R^2^ are preferred. c. Association significance (p-value) between alpha diversity (Shannon index) and batch/condition using Wilcoxon rank-sum test at α= 0.05. A non-significant association between alpha diversity and batch and a significant association between alpha diversity and condition are preferred. d. Cross-validated prediction accuracy (AUROC and F1) of the integrated microbiome data for the disease condition using random forest. Higher AUROC and F1 indicate a better prediction combining sensitivity and specificity.

For the IBD and CRC studies with relative abundance data, limma surprisingly amplified batch effects in the IBD data (the corresponding conditional signal was also amplified), with all other methods reduced batch effect as desired in both studies. ConQuR-rel, followed by MMUPHin and Percentile Normalization, were the most effective in mitigating batch variations. Also, limma and ConQuR-rel, followed by MMUPHin and ComBat, effectively retained the condition effects. Unfortunately, no methods were successful in removing the association between the richness/evenness of the microbiome and batch, except for Percentile Normalization in IBD. For the association between richness/evenness and condition, only ConQuR-rel and limma maintained or improved the condition signals in IBD and CRC data. Among the rest, ComBat and MMUPHin retained the key signal in the IBD study but failed to do so in the CRC study, while Harmony successfully improved the condition signal in the CRC study but not in the IBD study. Regarding the predictive ability, ConQuR-rel, followed by limma, ComBat, and MMUPHin, retained or even improved the condition signals.

Overall, in the real-world microbiome studies, we again conclude that different strategies thrive given different analytical emphases. However, our conclusions regarding batch removal and condition effect retention mostly align with those from simulations. The ConQuR family shows superior performance overall, while the ComBat family, limma, and MMUPHin can be dominating for different analytical goals. Notably, the ConQuR family is, again, slower than the rest of the strategies by runtime (Supplemental Figure 5d).

### MOSAIC-supported quantitative relevance evaluation for differentially abundant microbial markers from real-world microbiome studies

For real-world studies, FDR and sensitivity of differential abundance analysis cannot be evaluated without ground truths. We therefore devised a MOSAIC module to study the relevance of discovered biomarkers in relation to the diseases in literature (Methods). We follow the assumption that when a taxon and the disease are mentioned together in many publications, the taxon and the disease are strongly related. Therefore, we define a *strongly relevant* taxon as a taxon that has been reported along with the investigated disease and its related clinical terms in numerous publications (see *Relevance of real-world differentially abundant taxa via MOSAIC* of the Methods section for details). The numbers of *strongly relevant* taxa discovered in integrated data from each strategy in each of the four diseases are recorded in Supplemental Table 3. Consistently across diseases, merely any strongly relevant taxa were discovered in post-Percentile Normalization data (except for one taxa discovered in CRC), agreeing with simulation results. Other methods all retain similar number of or more taxa than without integration, showing the general power of integration strategies in retaining signal (Supplemental Table 4). Reassuringly, such discovered taxa also display a high degree of consensus.

We also explored the statistically significant differential taxa and the *strongly relevant* taxa among them (Supplemental Tables 3-4). For the autism study with taxonomic count data, ComBat-seq and MMUPHin identified *Clostridium XlVb*, the only discovered biomarker across all integration strategies which does turn out to be a *strongly relevant* taxon.

*Clostridium* has been extensively linked to autism and related disorders due to its production of exotoxins and propionate, which can worsen autism symptoms ^30,31^. For the CDI study with count data, there are 59 taxa discovered by one or more integration strategies, and all of them are *strongly relevant*. For example, *Akkermansia* and *Bacteroides* show a known protection effect against CDI infection ^32,33^. For the IBD study with relative abundance data, ComBat and Percentile Normalization are not sensitive, only identifying 2 and 0 *strongly relevant taxa*, while limma exhibits superior ability in extracting signals for IBD, which is followed by ConQuR-rel. Among the *strongly relevant* taxa that are detected, we highlight the discovery of 17 *Bacteroides* species – the decreased abundance of *Bacteroides* in IBD patients compared to healthy patients are well noted in literature ^34–36^. For CRC study with relative abundance data, we did not identify significant taxa in most integrated datasets, but 10 *strongly relevant* taxa are found from the ConQuR-rel integrated data, with one *strongly relevant* taxon discovered from the Percentile Normalization integrated data. This again elevates the ConQuR family’s ability to discover meaningful biological signals. These *strongly relevant* taxa identified by ConQuR-rel include species in *Streptococcus* and *Lactobacillus*, such as *S. milleri* and *L. delbrueckii*, and *L. plantarum. Streptococus* and *Lactobacillus* are some of the most known clinically relevant CRC adenoma biomarkers, the latter of which have been discussed to be used as probiotics for CRC management ^37,38^.

## Discussion

With the increasing prevalence of large-scale and consortium microbiome studies, there is a growing demand for rigorous microbiome data integration. Although many integration strategies have been borrowed or developed for microbiome data, a systematic pipeline to determine the best strategy is lacking. To address this gap, we develop MOSAIC, the first-of-its-kind pipeline that implements various integration approaches and compares their performance pertaining to various analysis aspects. We then used MOSAIC to comprehensively benchmark strategies on simulated and real-world datasets.

MOSAIC consists of three modules. The first module handles data-specific and strategy-specific pre-processing to ensure the best use. The second module applies these strategies to generate integrated data. The third module evaluates the effectiveness of the integration. This evaluation is based on the principle that a successful data integration strategy eliminates undesirable batch effects while preserving important biological signals. The specific metrics are inspired by important microbiome analyses – alpha and beta diversity analysis, microbiome-based predictive modelling, and differential abundance analysis – as well as runtime. The entire workflow is efficiently packaged by Snakemake ^26^, to ensure user friendliness. MOSAIC also supports a quantitative biomarker relevance evaluator. This tool provides a proxy for comparing the reliability of integration strategies in real-world differential analyses without ground truth. Additionally, it assesses how well each identified taxon aligns with historical literature on the disease. This helps researchers quickly narrow down microbes for further investigation.

Using MOSAIC, we compared different integration strategies across numerous simulated and real-world microbiome studies, encompassing various diseases, study sizes (sample size and microbial profile size), strengths and types of condition and batch effects, and numbers of data sources, including both 16S rRNA sequencing and shotgun metagenomic sequencing data, as well as taxonomic count and relative abundance data types. The results indicate that no single method excels in all aspects of analysis, but several general conclusions can be drawn. Percentile Normalization ^22^ achieves the most thorough batch effect removal but also eliminates the condition effect. ComBat ^17^ (for relative abundance), limma ^19^, and Harmony ^39^ result in partial batch removal and slight reduction in condition signals. MMUPHin ^21^ excels at maintaining or even enhancing the condition effect while moderately eliminating batch effects. ComBat-seq ^18^ (for taxonomic count) and the ConQuR family ^20^ are generally superior in both batch effect removal and condition effect retention.

Our study also comes with limitations. Firstly, with the ever-growing number of microbiome data integration strategies, our study might not cover all existing strategies. However, MOSAIC’s flexible design enables easy inclusion of future strategies. Validating strategies on real-world data is also challenging, as comparing sensitivity and FDR in differential abundance analyses is difficult without known ground truth taxa. Currently, our approach relies on user-defined clinical terms to identify publications linking taxa to specific diseases, which may result in potential double-counting when multiple terms appear in a single publication. To mitigate this, we defined distinct, broad clinical terms to cover various research areas relevant to each disease. We recommend that users carefully select clinical terms to ensure unbiased and comprehensive coverage of relevant research.

Overall, MOSAIC helps microbiome researchers focus on downstream analyses with minimal concern about data integration in the era of big microbiome data studies. Furthermore, the MOSAIC-powered integration strategy benchmarks provide valuable references for researchers in the field. Enabled by its well-designed and flexible structure, MOSAIC opens new avenues for further development. First, more metrics evaluating data integration effectiveness can be incorporated to meet the increasing need for relevant analytical aspects, such as network analysis, mediation, or even causal analysis. Next, while the MOSAIC-supported real-world biomarker relevance evaluator identifies the historical relevance of a biomarker, experimental investigations on novel markers are necessary to corroborate their relevance. As microbiome studies continue to grow in size and biomedical importance, the MOSAIC pipeline, which supports benchmarking across various integration strategies by covering a broader spectrum of analyses, will become increasingly well-regarded among biomedical and methodological investigators within the field.

## Methods

### Details of MOSAIC

Our system streamlines the implementation and evaluation of microbiome data integration strategies on data sets of interest. The pipeline supports a wide range of integration tools, and facilitates a flexible, comprehensive, and fair comparison. Based on the automatically generated final report summarizing a broad spectrum of evaluation metrics along with the accompanied visualizations, the user can choose the optimal approach to address their specific research need (Figure 1). Implementation and usage details of MOSAIC are on GitHub (https://github.com/tommyfuu/MOSAIC).

#### Module 1. Data Pre-Processing

Module 1 loads in and pre-processes the microbiome data. In terms of data loading, MOSAIC has wrappers that cater to two types of data inputs – in .csv file format (see the example on GitHub, from the MicrobiomeHD database ^24^) or R objects, including plain types such as matrix/data frame and the special type for microbiome data, the phyloseq object (specifically from the CuratedMetagenomicData platform to acquire real world data ^29,40^).

Then, MOSAIC conducts generic pre-processing on the input microbiome and meta data. This includes taxonomic aggregation – agglomerated to genus level for 16S rRNA sequencing data and to species level for shotgun metagenomics sequencing data, zero lineage filtering, and low-quality sample removal – filtered out the samples with library sizes below a particular percentile of all the samples, which can be specified by the user such as 5%. Note that all filtering is conducted with the R package phyloseq ^37^. MOSAIC also generates descriptive statistics of the processed data, including the number of samples in each condition and in each batch as well as the total number taxa (see the Supplemental Table 2, for four real-world data investigated in the paper).

#### Module 2. Data Integration

Module 2 applies various embedded data integration strategies to the pre-processed data. The pipeline offers several options: ComBat ^17^, ComBat-seq ^18^, limma ^19^, MMUPHin ^21^, the ConQuR family ^20^, Harmony ^39^, and Percentile Normalization ^22^ as potential data integration strategies. Among them, ComBat, ComBat-seq, and MMUPHin all employ the empirical Bayes framework within their respective models – Gaussian linear regression, negative binomial regression, and an enhanced Gaussian linear regression that includes a component to address the zero-inflated nature of microbiome data. limma uses frequentist linear regression to explicitly model and remove batch effects. Harmony is a similarity-based methods to align similar samples to a common distribution in a reduced latent space, regardless of batch. Nonparametric strategies that do not assume any parametric distributions, the ConQuR family and Percentile Normalization, are also included. The former match the conditional distribution of microbiome data via zero-inflated quantile regression, and the latter uses the marginal percentile matching principle to even out the differences across batches. These approaches make distinct assumptions on the underlying data and may work only on taxonomic count or relative abundance. Please refer to Supplemental Table 1 for a summary.

Users can select a subset, or all the strategies based on the data type of interest (either taxonomic count or relative abundance). Module 2 conducts strategy-specific normalization to ensure the best use of the integration methods. For each selected strategy, the generic processed data from Module 1 is further normalized according to the strategy’s requirements (See Supplemental Table 2). Finally, the normalized data is integrated by the corresponding strategy to generate a homogeneous data set.

#### Module 3. Result Evaluation and visualization

The final and most important module evaluates the effectiveness of the data integration methods. It provides a comprehensive report and visual displays of the integrated data by different strategies, based on which the investigators can choose a strategy and the corresponding cleaned data for downstream analyses.

As the overarching goal of data integration is to remove batch effects while maintaining condition effects, we designed four types of metrics covering the most common microbiome analyses to ensure a comprehensive comparison (Figure 1, part 3).

##### Alpha diversity

reflects the richness and/or evenness of microbial communities. Successful data integration is indicated by a weaker association between alpha diversity and batch, alongside a stable or stronger association with the condition. The Shannon index, the most widely used alpha diversity metric, is computed using the scikit-bio Python package ^40^. Associations between the Shannon index and batch or condition are then assessed via the Wilcoxon rank-sum test (for two groups) or the Kruskal–Wallis test (for three or more groups) using the statsmodels Python package ^41^. In simulation studies, the proportion of p-values below the significance level α represents type I error for batch effects (where no batch effect is expected) and power for the condition (where a pronounced effect is expected). An ideal integration method controls type I error at α while maximizing statistical power. For real-world data analysis, where ground truth and large numbers of replicates are not available, a p-value greater than α for batch suggests no batch effect, while a p-value less than α for the condition indicates a significant condition effect.

##### Beta diversity

measures between-sample distances to characterize differences in microbiome compositions. To quantify how much variability in the microbiome data (captured by beta diversity) is explained by a particular variable (batch or condition), we use PERMANOVA ^25^ R^2^. Successful data integration is reflected by a decrease of batch R^2^ and a stable or increased condition R^2^. For count data, PERMANOVA R^2^ is calculated using both Bray-Curtis and Aitchison distances, while only Bray-Curtis is applied to relative abundance data. In simulation studies, the average batch R^2^ and condition R^2^ are computed for an accurate estimates of batch effects and condition effects. In real-world analysis, we simply rely on the one-time estimates.

Machine learning models have been increasingly used to predict disease phenotypes based on the host microbiome for diagnostic purposes ^42,43^. The ***microbiome-based prediction accuracy*** of condition can reflect how well the microbiome explains the disease condition. MOSAIC leverages the accuracy of random forest model as another measure of condition effect retention (in the reverse direction of condition R^2^, which indicates how well the condition explains the microbiome). We train a random forest model (with 100 estimators, a maximum depth of 2, and random seed = 0) using 5-fold cross validation, and compute the average model accuracy, AUC scores, macro and weighted precisions, macro and weighted recalls, and macro and weighted F1 scores, using the scikit-learn python package ^44^. Higher values for these metrics indicate better integration.

A recurring objective of microbiome studies is to identify individual taxa that are associated with the disease condition through ***differential abundance analysis***. Successful integration preserves or enhances the ability to discover differentially abundant taxa while minimizing false discoveries. In simulation studies, where the ground truth is known, MOSAIC evaluates the integration strategies by calculating the average sensitivity and false discovery rate (FDR) across simulation rounds. Specifically, for each round, we use linear regression model with Benjamini-Hochberg correction to compute FDR-adjusted p-values, using the statsmodels python package ^41^. The best integration method is indicated by a controlled FDR (at α=0.05) and maximized sensitivity. For real-world analysis, where the ground truth is unknown, the relevance of identified taxa is assessed using existing literature, as detailed in the next subsection.

At last, in addition to numerical tables that summarize the metrics above, MOSAIC provides visualizations for each metric for a straightforward comparison, such as boxplots for alpha diversity, principal coordinates analysis (PCoA) plot for beta diversity, performance curves for the predictive model, and the sensitivity/FDR curves across various scenarios for differential abundance analysis in simulation studies.

For our benchmarking work, all simulation, integration strategies, and evaluations were run with 2 CPUs with 20G memory on a distributed computing system.

### Relevance of real-world differentially abundant taxa via MOSAIC

In real-world differential abundance analysis, sensitivity and FDR cannot be quantitatively assessed due to the lack of ground truth. To address this, we propose using previously published literature, accessible via the PubTator literature search engine ^45^, to evaluate the relevance of the taxa identified from the integrated data.

The approach is semi-automated. Given a condition, the users, in collaboration with domain experts, compile a short list of key terms relevant to that condition. For the four diseases in our real-world analysis, we generated the following key terms:

- Autism: autism, autism spectrum disorders, gut-brain axis, neurological disorders, nervous systems
- CDI: *Clostridioides* difficile, diarrhea, antibiotics, toxins, infection
- IBD: Irritable Bowel Syndrome, Crohn’s disease, FODMAP diet, abdominal pain, inflammation
- CRC: colon cancer, colorectal cancer, diarrhea, rectal bleeding, MSI subtype

Next, using PubTator, we assess how many previously published papers have discussed the relationship between each identified taxon and each of the key terms for the condition. The search is implemented in scale through a Python web-mining script that automatically queries PubTator with “BIOMARKER_NAME” and “CLINICAL TERM” and returns the number of publications found.

For each taxon, we count how many of the five key terms yield at least one relevant publication, and how many yield at least ten. A taxon is considered *strongly relevant taxon* if it returns at least one publication for all five terms, or at least ten publications for three of the five terms. For each real-world study, we then count the number of *strongly relevant taxa* identified from the integrated microbiome data generated by the various strategies. The strategy that identifies the most *strongly relevant taxa* is considered the most effective at retaining or improving useful signals.

### Simulation settings

To evaluate the integration strategies using simulated data, we utilized the MIDASim^27^ microbiome data simulator, which generates a microbiome data based on specified requirements while mimicking a real data set. Specifically, MIDASim has a parametric version and a non-parametric version, and here we used the nonparametric version because it mimics real data better than the parametric version. We used a gut microbiome data set from the Inflammatory Bowel Disease Multi-omics Database (IBDMDB) project, which is a template data of the Integrative Human Microbiome Project’s database^5,28,46^. We aggregated the data to the genus level, removed samples with library size below 2000, and filtered out taxa present in less than 5% of the samples, resulting in 301 taxa and 150 samples. Our aim was to simulate alike microbiome data with 301 taxa and 450 samples.

We simulated 2 conditions (*condition* 1 vs. 0) and 2 batches (*batch* 1 vs. 0) for the 450 samples from a joint Bernoulli distribution with p_condition_ = 0.5, p_Batch_ = 0.5, and odds ratio (OR) = 1, 1.25, or 1.5. OR=1 indicates batch and condition are independent, while OR ≠ 1 indicates batch and condition are correlated and batch can be a potential confounder of the condition-microbiome relationships. We further considered the scenarios that library size is library size is introduced as part of the batch effect, by setting p_batch=1, i_ = 1/(1+exp(*libsize*_i_) instead of 0.5 for sample *i*. We chose 50 taxa to be differentially abundant between Condition 1 and 0. To avoid bias from manual selection, we ordered the taxa according to their average abundances and selected the taxa from the least abundant to most abundant evenly across the spectrum of abundance. Also, we assumed that all 301 taxa would be affected by batch.

We first fitted MIDASim to the processed IBD data to obtain *v*_*0*_, which is the taxon-wise mean value vector in the starting data. Next, we modified the parameter to introduce condition and batch effects. Specifically, for the 50 differentially abundant taxa, we randomly permuted the corresponding means in *v*_*0*_ to obtain *v*_*C*_, which enables the additive effect of *condition* 1 while keeping the total sum of condition effects remaining the same. We did similar adjustment for taxa from samples belonging to *batch* 1, with *v*_*B*_ obtained from permuting *v*_*0*_ as well.

For each round’s simulated microbiome data containing 301 taxa and 450 samples, the 450 samples’ library sizes were randomly sampled with replacement from the 150 starting samples. Then, we considered a wide range of combinations of the condition and batch effects. The pool of condition effect is *s*_*C*_ is in {0.25, 0.5, 0.75, 1} and the pool of batch effect is *s*_*B*_ ={0, 0.25, 0.5, 0.75, 1}. These combinations comprehensively encapsulated three scenarios:

(1) batch effect *s*_*B*_ > condition effect *s*_*C*_
(2) batch effect *s*_*B*_ = condition effect *s*_*C*_
(3) batch effect *s*_*B*_ < condition effect *s*_*C*_ (including no batch effect)

Given *v*_*0*_, we then fed in MIDASim with a taxon-wise mean value vector v_i_ = (1-*s*_*B*_ *batch*_i_-*s*_*C*_*condition*_i_)*v*_*0*_+*s*_*B*_ *batch*_i_ *v*_*B*_*+s*_*C*_ *condition*_i_ *v*_*C*_ to modify the starting data. Note that due to this constraint, the sum of *s*_*B*_ and *s*_*C*_in a data set cannot exceed 1.

For each setting based on condition/batch effect, odds ratio, whether library size is part of batch effect, we repeated the simulation 1,000 rounds. The performances of the integration strategies were summarized over the 1,000 trials.

## Reference

1. Xie, H. et al. Shotgun Metagenomics of 250 Adult Twins Reveals Genetic and Environmental Impacts on the Gut Microbiome. Cell Syst. 3, 572-584.e3 (2016).

2. Zhou, W. et al. Longitudinal multi-omics of host-microbe dynamics in prediabetes. Nature 569, 663–671 (2019).

3. Hou, K. et al. Microbiota in health and diseases. Signal Transduct. Target. Ther. 7, 135 (2022).

4. Schirmer, M. et al. Dynamics of metatranscription in the inflammatory bowel disease gut microbiome. Nat. Microbiol. 3, 337–346 (2018).

5. IBDMDB Investigators et al. Multi-omics of the gut microbial ecosystem in inflammatory bowel diseases. Nature 569, 655–662 (2019).

6. Li, J. et al. An integrated catalog of reference genes in the human gut microbiome. Nat. Biotechnol. 32, 834–841 (2014).

7. Nielsen, H. B. et al. Identification and assembly of genomes and genetic elements in complex metagenomic samples without using reference genomes. Nat. Biotechnol. 32, 822–828 (2014).

8. Feng, Q. et al. Gut microbiome development along the colorectal adenoma–carcinoma sequence. Nat. Commun. 6, 6528 (2015).

9. Hannigan, G. D., Duhaime, M. B., Ruffin, M. T., Koumpouras, C. C. & Schloss, P. D. Diagnostic Potential and Interactive Dynamics of the Colorectal Cancer Virome. mBio 9, e02248–18 (2018).

10. Thomas, A. M. et al. Metagenomic analysis of colorectal cancer datasets identifies cross-cohort microbial diagnostic signatures and a link with choline degradation. Nat. Med. 25, 667–678 (2019).

11. Yachida, S. et al. Metagenomic and metabolomic analyses reveal distinct stage-specific phenotypes of the gut microbiota in colorectal cancer. Nat. Med. 25, 968–976 (2019).

12. The Human Microbiome Project Consortium. Structure, function and diversity of the healthy human microbiome. Nature 486, 207–214 (2012).

13. Danko, D. et al. A global metagenomic map of urban microbiomes and antimicrobial resistance. Cell 184, 3376-3393.e17 (2021).

14. Department of Clinical and Toxicological Analyses, School of Pharmaceutical Sciences, University of Sao Paulo, Sao Paulo, Brazil et al. Integrative Biology Approaches Applied to Human Diseases. in Computational Biology (eds. Division of Biomedical Science, University of the Highlands and Islands, U. & Husi, H.) 19–36 (Codon Publications, 2019). doi:10.15586/computationalbiology.2019.ch2.

15. Leek, J. T. et al. Tackling the widespread and critical impact of batch effects in high-throughput data. Nat. Rev. Genet. 11, 733–739 (2010).

16. Ioannidis, J. P. A., Patsopoulos, N. A. & Evangelou, E. Heterogeneity in Meta-Analyses of Genome-Wide Association Investigations. PLoS ONE 2, e841 (2007).

17. Johnson, W. E., Li, C. & Rabinovic, A. Adjusting batch effects in microarray expression data using empirical Bayes methods. Biostatistics 8, 118–127 (2007).

18. Zhang, Y., Parmigiani, G. & Johnson, W. E. ComBat-seq: batch effect adjustment for RNA-seq count data. NAR Genomics Bioinforma. 2, lqaa078 (2020).

19. Ritchie, M. E. et al. limma powers differential expression analyses for RNA-sequencing and microarray studies. Nucleic Acids Res. 43, e47–e47 (2015).

20. Ling, W. et al. Batch effects removal for microbiome data via conditional quantile regression. Nat. Commun. 13, 5418 (2022).

21. Ma, S. et al. Population structure discovery in meta-analyzed microbial communities and inflammatory bowel disease using MMUPHin. Genome Biol. 23, 208 (2022).

22. Gibbons, S. M., Duvallet, C. & Alm, E. J. Correcting for batch effects in case-control microbiome studies. PLOS Comput. Biol. 14, e1006102 (2018).

23. Xiao, L., Zhang, F. & Zhao, F. Large-scale microbiome data integration enables robust biomarker identification. Nat. Comput. Sci. 2, 307–316 (2022).

24. Duvallet, C., Gibbons, S. M., Gurry, T., Irizarry, R. A. & Alm, E. J. Meta-analysis of gut microbiome studies identifies disease-specific and shared responses. Nat. Commun. 8, 1784 (2017).

25. Anderson, M. J. Permutational Multivariate Analysis of Variance (PERMANOVA). in Wiley StatsRef: Statistics Reference Online 1–15 (John Wiley & Sons, Ltd, 2017). doi:10.1002/9781118445112.stat07841.

26. Köster, J. & Rahmann, S. Snakemake—a scalable bioinformatics workflow engine. Bioinformatics 28, 2520–2522 (2012).

27. He, M., Zhao, N. & Satten, G. A. MIDASim: a fast and simple simulator for realistic microbiome data. Microbiome 12, 135 (2024).

28. HMP2Data: 16s rRNA sequencing data from the Human Microbiome Project 2.

29. Pasolli, E. et al. Accessible, curated metagenomic data through ExperimentHub. Nat. Methods 14, 1023–1024 (2017).

30. Kang, D.-W. et al. Reduced Incidence of Prevotella and Other Fermenters in Intestinal Microflora of Autistic Children. PLOS ONE 8, e68322 (2013).

31. Argou-Cardozo, I. & Zeidán-Chuliá, F. Clostridium Bacteria and Autism Spectrum Conditions: A Systematic Review and Hypothetical Contribution of Environmental Glyphosate Levels. Med. Sci. 6, 29 (2018).

32. Deng, H. et al. Bacteroides fragilis Prevents Clostridium difficile Infection in a Mouse Model by Restoring Gut Barrier and Microbiome Regulation. Front. Microbiol. 9, (2018).

33. Wu, Z. et al. Akkermansia muciniphila Ameliorates Clostridioides difficile Infection in Mice by Modulating the Intestinal Microbiome and Metabolites. Front. Microbiol. 13, (2022).

34. Quaglio, A. E. V., Grillo, T. G., Oliveira, E. C. S. D., Stasi, L. C. D. & Sassaki, L. Y. Gut microbiota, inflammatory bowel disease and colorectal cancer. World J. Gastroenterol. 28, 4053–4060 (2022).

35. Yao, C. et al. Significant Differences in Gut Microbiota Between Irritable Bowel Syndrome with Diarrhea and Healthy Controls in Southwest China. Dig. Dis. Sci. 68, 106– 127 (2023).

36. Sundin, J., Rangel, I., Repsilber, D. & Brummer, R.-J. Cytokine Response after Stimulation with Key Commensal Bacteria Differ in Post-Infectious Irritable Bowel Syndrome (PI-IBS) Patients Compared to Healthy Controls. PLOS ONE 10, e0134836 (2015).

37. Agnes, A. et al. Association between colorectal cancer and Streptococcus gallolyticus subsp. pasteuranus (former S. bovis) endocarditis: clinical relevance and cues for microbiota science. Case report and review of the literature. Eur. Rev. Med. Pharmacol. Sci. 25, 480–486 (2021).

38. Sivamaruthi, B. S., Kesika, P. & Chaiyasut, C. The Role of Probiotics in Colorectal Cancer Management. Evid. Based Complement. Alternat. Med. 2020, e3535982 (2020).

39. Korsunsky, I. et al. Fast, sensitive and accurate integration of single-cell data with Harmony. Nat. Methods 16, 1289–1296 (2019).

40. McMurdie, P. J. & Holmes, S. phyloseq: An R Package for Reproducible Interactive Analysis and Graphics of Microbiome Census Data. PLOS ONE 8, e61217 (2013).

41. Seabold, S. & Perktold, J. Statsmodels: Econometric and Statistical Modeling with Python. in 92–96 (Austin, Texas, 2010). doi:10.25080/Majora-92bf1922-011.

42. Su, Q. et al. Faecal microbiome-based machine learning for multi-class disease diagnosis. Nat. Commun. 13, 6818 (2022).

43. Marcos-Zambrano, L. J. et al. Applications of Machine Learning in Human Microbiome Studies: A Review on Feature Selection, Biomarker Identification, Disease Prediction and Treatment. Front. Microbiol. 12, (2021).

44. Pedregosa, F. et al. Scikit-learn: Machine Learning in Python. Mach. Learn. PYTHON.

45. Wei, C.-H., Allot, A., Leaman, R. & Lu, Z. PubTator central: automated concept annotation for biomedical full text articles. Nucleic Acids Res. 47, W587–W593 (2019).

46. Proctor, L. M. et al. The Integrative Human Microbiome Project. Nature 569, 641– 648 (2019).

