## Supplementary material for "MOSAIC: A Pipeline for MicrobiOme Studies Analytical Integration and Correction": Supp 1

**Further descriptions regarding simulation results**

Evaluation results based on simulated taxonomic count microbiome data remain mostly consistent with Figure 2 and the Results section when varying settings with different levels of confounding effects between batch and condition and varied confounding statuses between batch and library size (Supplemental Figures 1-2).

On simulated relative abundance data, the comparison results were mostly replicated (Supplemental Figures 3-4). Percentile Normalization thoroughly removed both batch and condition effects. ConQuR-rel demonstrated superior performance overall, effectively mitigated batch effects while retaining or even improving the condition effects. ComBat was the runner-up. However, its effectiveness in correcting for batch was not as pronounced as its extension for count data, ComBat-seq, due to its Gaussian assumption being far from the zero-inflated relative abundance data. Harmony, limma, and MMUPHin maintained the condition effect, while they could only reduce partial batch variation. For association testing, ConQuR-rel, MMUPHin, and Percentile Normalization were the only strategies that consistently controlled FDR below 0.05. ComBat, limma, especially Harmony exhibited inflated FDR. Among those that controlled FDR, only ConQuR-rel and MMUPHin improved sensitivity.

**Supplemental Figure 1. Evaluation of data integration strategies on simulated microbiome data.** Data type is count and confounding between batch effect and library sizes exist. Odds ratios between batch and condition for the two panels are 1, 1.25. Other simulation settings are the same as in Figure 2.

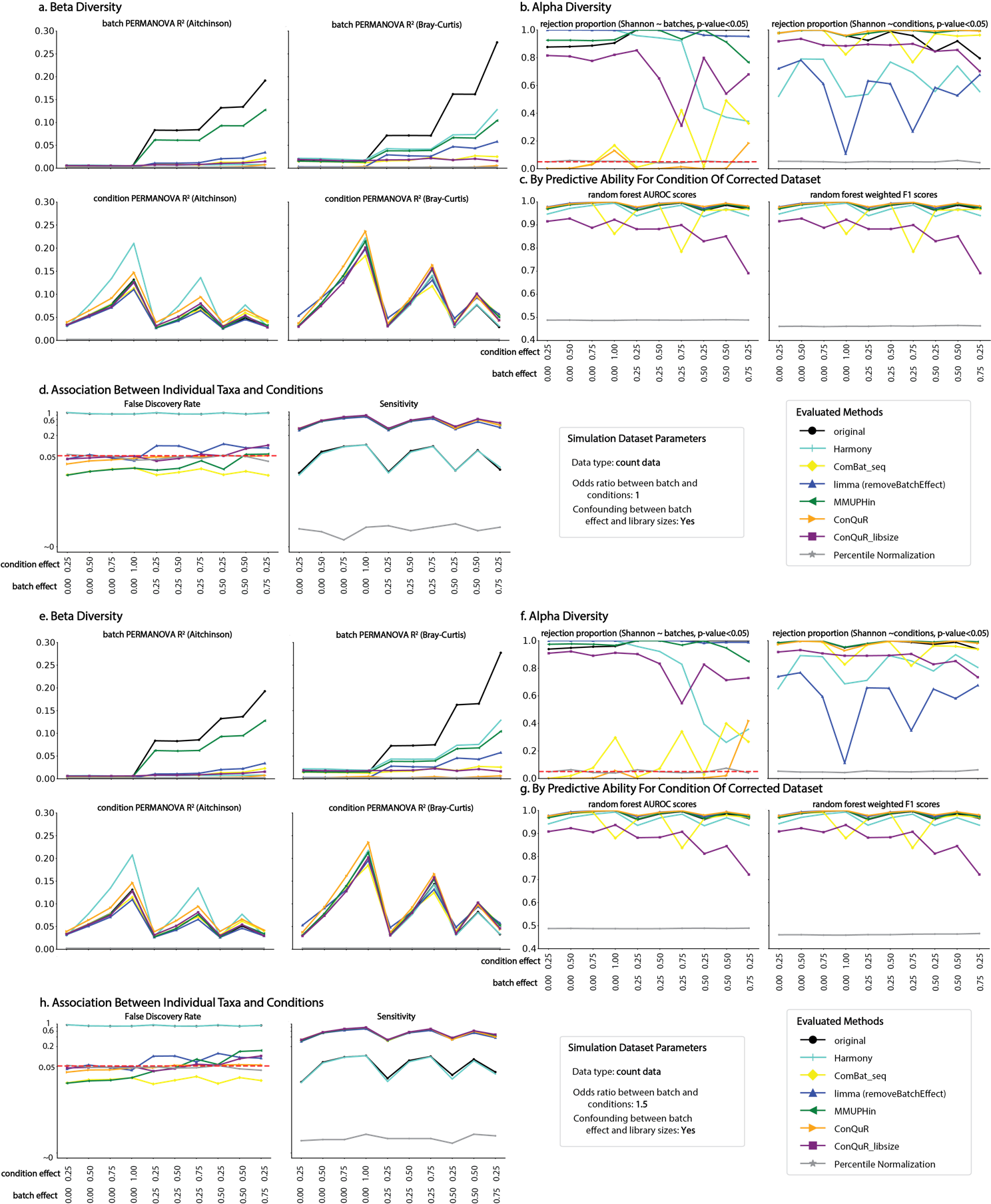

**Supplemental Figure 2. Evaluation of data integration strategies on simulated microbiome data.** Data type is count and confounding between batch effect and library sizes does not exist. Odds ratios between batch and condition for the two panels are 1, 1.25, 1.5. Other simulation settings are the same as in Figure 2.

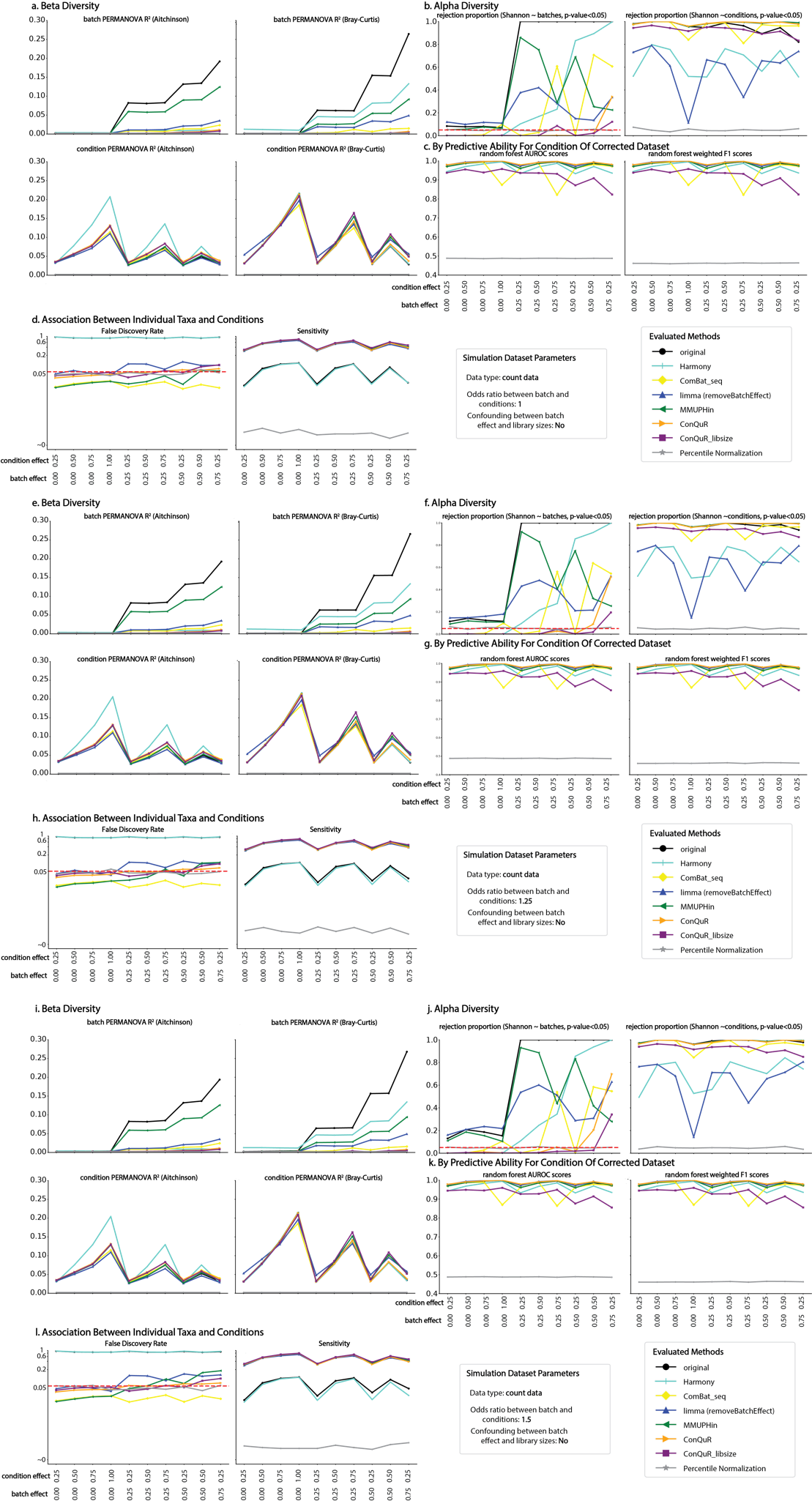

**Supplemental Figure 3. Evaluation of data integration strategies on simulated microbiome data.** Data type is relative abundance and confounding between batch effect and library sizes exists. Odds ratios between batch and condition for the two panels are 1, 1.25, 1.5. Other simulation settings are the same as in Figure 2.

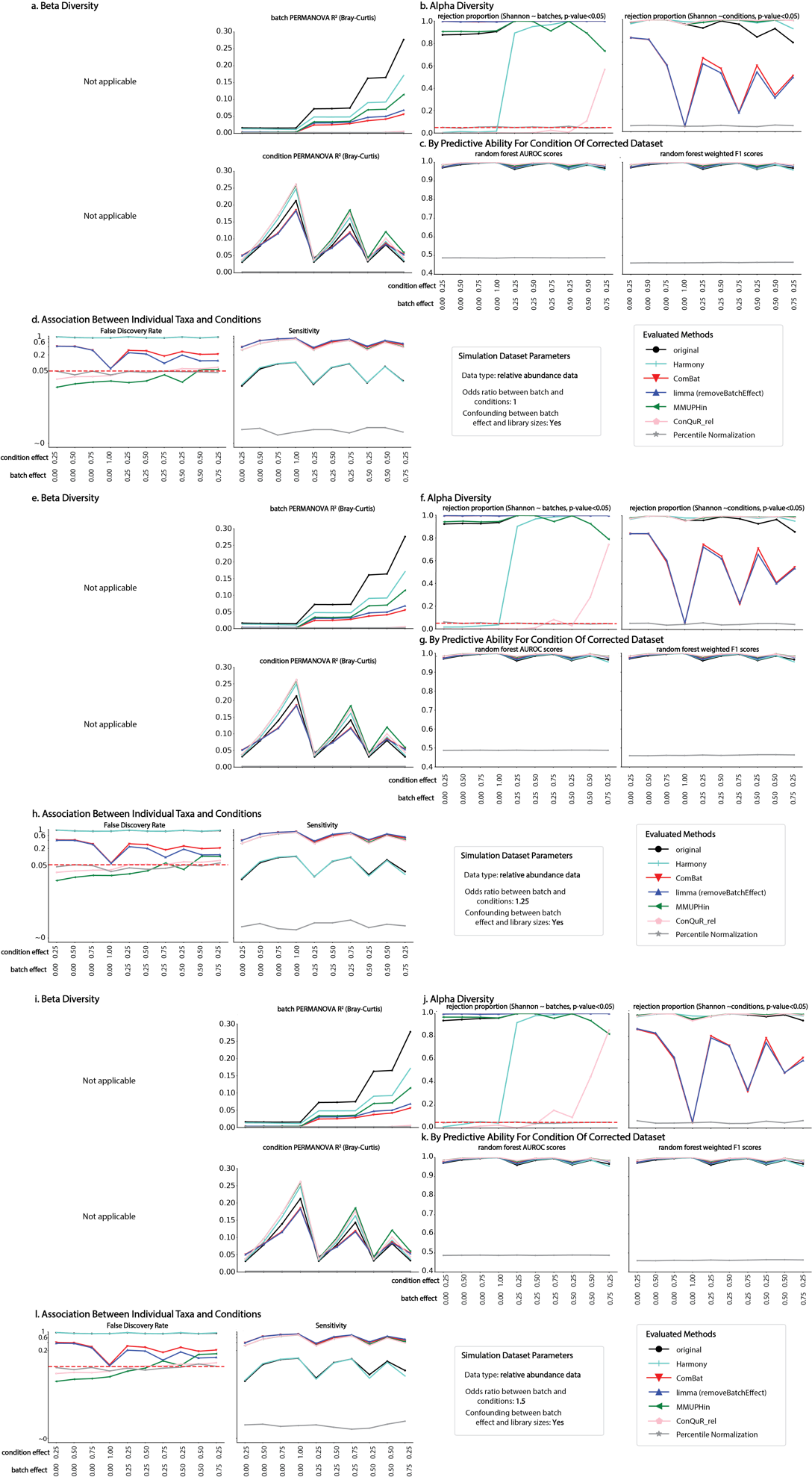

**Supplemental Figure 4. Evaluation of data integration strategies on simulated microbiome data.** Data type is relative abundance and confounding between batch effect and library sizes does not exist. Odds ratios between batch and condition for the two panels are 1, 1.25, 1.5. Other simulation settings are the same as in Figure 2.

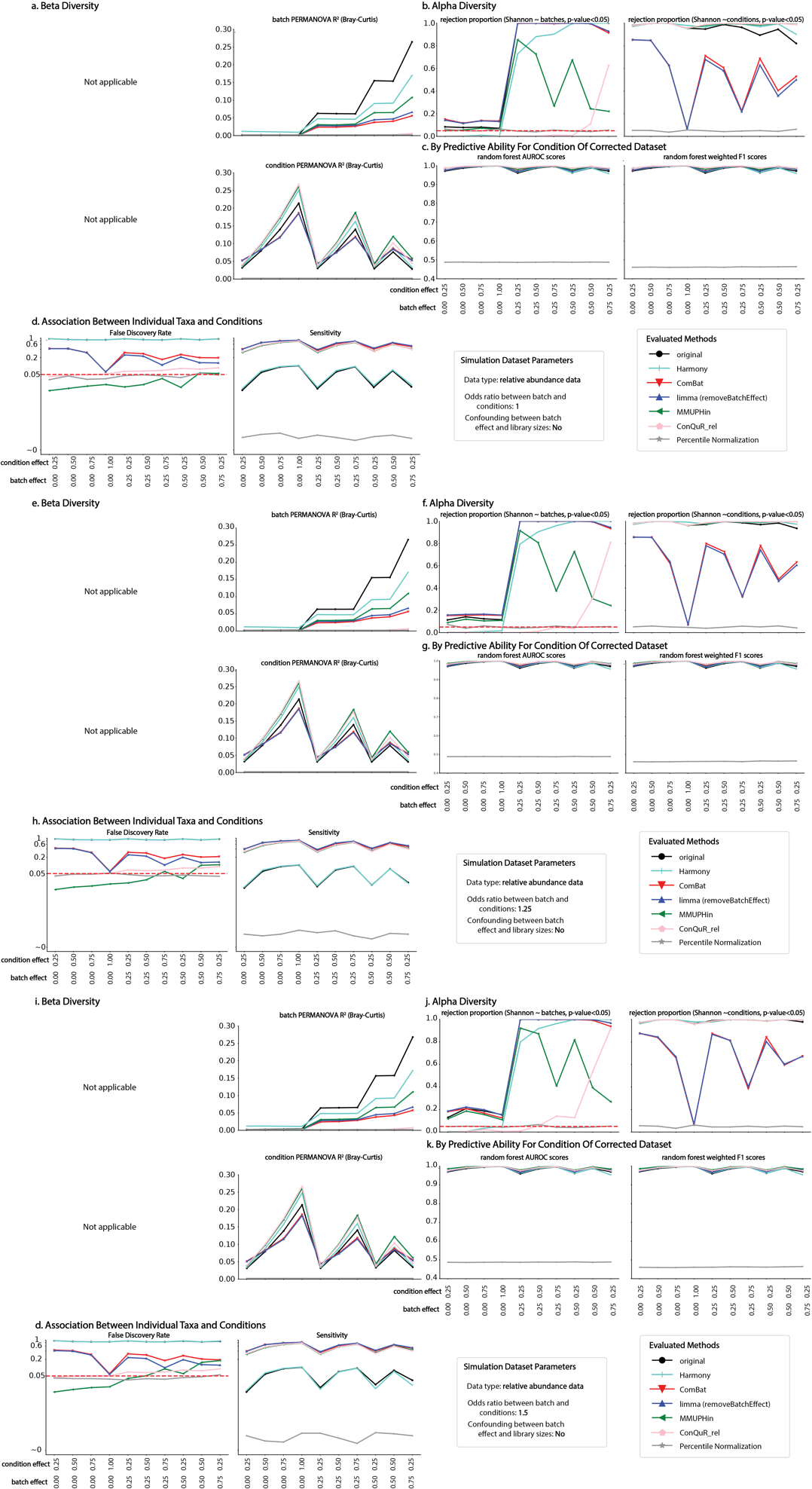
**Supplemental Figure 5. Runtime comparison of data integration strategies on simulated microbiome data.** Runtime (in seconds) are visualized in log2 scale.

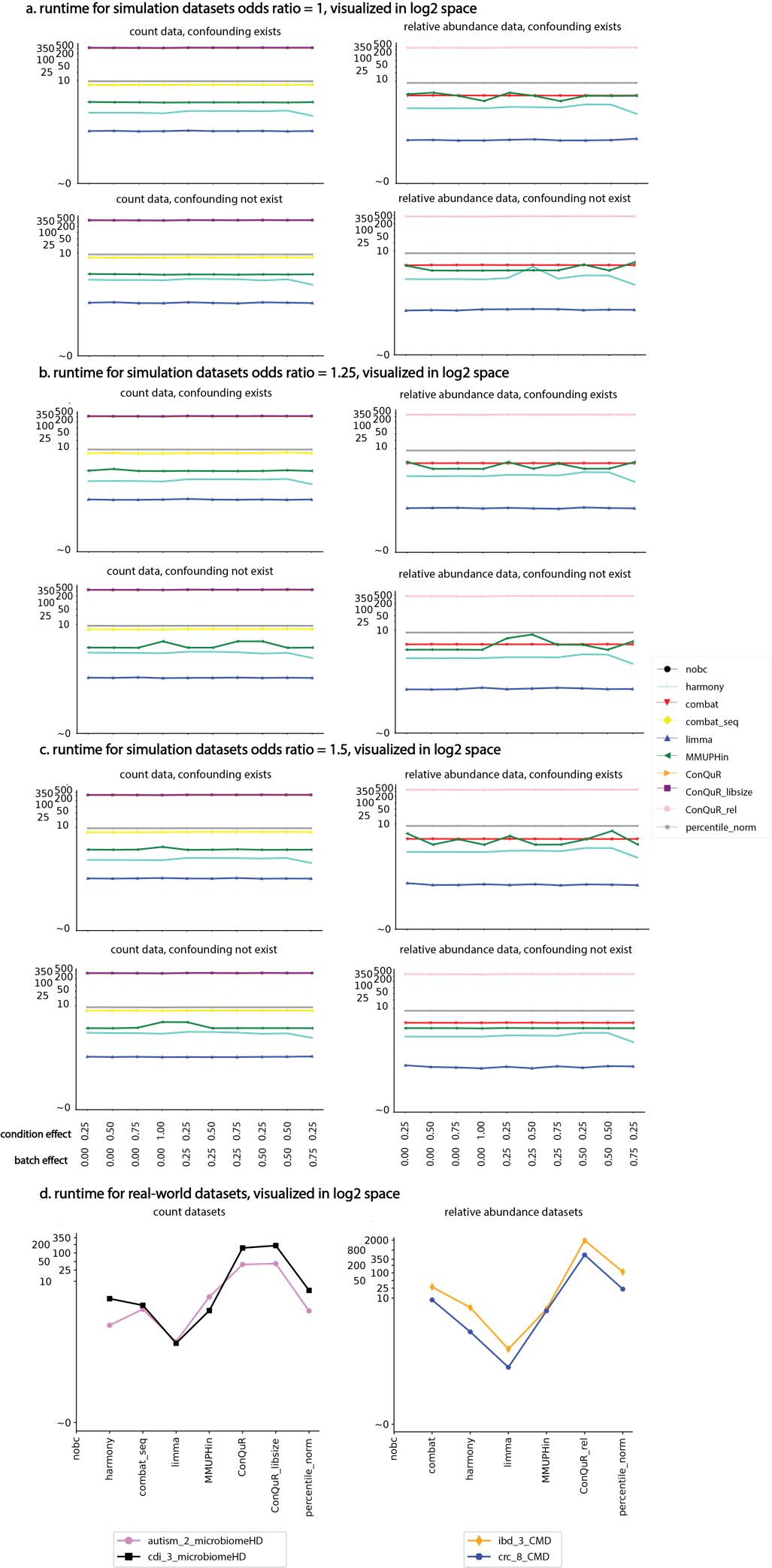

**Supplemental Table 1. Data integration strategies for microbiome data evaluated in this study.**

| Methods | For taxonomic counts | For relative abundance | Pre-processing requirements |
| --- | --- | --- | --- |
| *Combat_Seq* ^1^ | Yes | Yes | NA |
| *ComBat* ^2^ | No | Yes | clr |
| *limma* ^3^ | Yes | Yes | clr |
| *harmony* ^4^ | Yes | Yes | NA |
| *ConQuR* ^5^ | Yes | No | NA |
| *ConQuR_libsize* ^5^ | Yes | No | NA |
| *ConQuR_rel* ^5^ | No | Yes | NA |
| *MMUPHin* ^6^ | Yes | Yes | NA |
| *Percentile Normalization* ^7^ | Yes | Yes | Requires case-controlled data |

**Supplemental Table 2. Summary of processed real-world data evaluated in this study.**

| Dataset name | # samples* # taxa | batches | Condition/# samples |
| --- | --- | --- | --- |
| autism_MicrobiomeHD | 134*71 | Son et al. ^8^ | ASD/59; Healthy/44 |
|  |  | Kang et al. ^9^ | ASD/15; Healthy/16 |
| CDI_MicrobiomeHD | 461*94 | Schubert et al. ^10^ | CDI/83; Healthy/154; Diarrheal wo CDI/79 |
|  |  | Vincent et al. ^11^ | CDI/22; Healthy/23 |
|  |  | Youngster et al. ^12^ | CDI/27; Healthy/18; CDI post FMT Treatment/55 |
| IBD_CMD | 2021*597 | HMP ^13,14^ | IBD/1138; Healthy/414. |
|  |  | Li et al. ^15^ | IBD/45; Healthy/0. |
|  |  | Nielsen et al. ^16^ | IBD/140; Healthy/229. |
| CRC_CMD | 497*576 | Feng et al. ^17^ | CRC/8; Healthy/16. |
|  |  | Hannigan et al. ^18^ | CRC/23; Healthy/18. |
|  |  | Thomas et al. ^19^ | CRC/15; Healthy/13. |
|  |  | Yachida et al. ^20^ | CRC/61; Healthy/236. |
|  |  | Zeller et al. ^21^ | CRC/41; Healthy/56. |

**Supplemental Table 3. Number of *strongly relevant taxa* discovered from integrated dataset of each strategy in each evaluated real-world study.**

| Dataset name | Integration strategies | Number of strongly relevant taxa discovered |
| --- | --- | --- |
| autism_MicrobiomeHD | *No integration* | 0 |
|  | *Harmony* | 0 |
|  | *ComBat-seq* | 1 |
|  | *limma* | 0 |
|  | *MMUPHin* | 1 |
|  | *ConQuR* | 0 |
|  | *ConQuR-libsize* | 0 |
|  | *Percentile Normalization* | 0 |
| CDI_MicrobiomeHD | *No integration* | 31 |
|  | *Harmony* | 32 |
|  | *ComBat-seq* | 25 |
|  | *limma* | 37 |
|  | *MMUPHin* | 31 |
|  | *ConQuR* | 36 |
|  | *ConQuR-libsize* | 26 |
|  | *Percentile Normalization* | 0 |
| IBD_CMD | *No integration* | 48 |
|  | *Harmony* | 51 |
|  | *ComBat-seq* | 2 |
|  | *limma* | 181 |
|  | *MMUPHin* | 53 |
|  | *ConQuR-rel* | 74 |
|  | *Percentile Normalization* | 0 |
| CRC_CMD | *No integration* | 0 |
|  | *Harmony* | 0 |
|  | *ComBat* | 0 |
|  | *limma* | 0 |
|  | *MMUPHin* | 0 |
|  | *ConQuR-rel* | 10 |
|  | *Percentile Normalization* | 1 |
